# Convergent Evolution of A-Lineage (Clade 19B) SARS-CoV-2 Spike Sequences with B-Lineage Variants of Concern Affects Virus Replication in a Temperature-Dependent Manner on Human Nasal Epithelial Cell Cultures

**DOI:** 10.1101/2023.03.03.531067

**Authors:** Steve Yoon, Eddy Anaya, Jaiprasath Sachithanandham, Benjamin Pinsky, David Sullivan, Heba H. Mostafa, Andrew Pekosz

## Abstract

The first three months of the COVID-19 pandemic was dominated by two SARS-CoV-2 lineages: A-lineages (Clade 19B) and B-lineages (Clade 19A). However, with the emergence of the Spike D614G substitution in B.1 lineages (Clade 20A), both early lineages were outcompeted and remained near-extinction from mid-2020 onwards. In early-2021, there was a re-emergence and persistence of novel A-lineage variants with substitutions in the Spike gene resembling those found in Variants of Concern (VOCs). An early A.3 variant (MD-HP00076/2020) and three A.2.5 variants (MD-HP02153/2021, MD-HP05922/2021 and CA-VRLC091/2021) were isolated and characterized for their genomic sequences, antibody neutralization, and *in vitro* replication. All A.2.5 isolates had five Spike mutations relative to the A.3 variant sequence: D614G, L452R, Δ141-143, D215A, and ins215AGY. Plaque reduction neutralization assays demonstrated that A.2.5 isolates had a 2.5 to 5-fold reduction in neutralization using contemporaneous COVID-19 convalescent plasma when compared to A.3. *In vitro* viral characterization in VeroE6 cell lines revealed that the A.3 isolate grew faster and spread more than A.2.5. On VeroE6-TMPRSS2 cells, significant syncytia formation was also observed with the A.2.5 isolates, however Spike cleavage efficiency did not explain these differences. In human nasal epithelial cell (hNEC) cultures, the A.2.5 isolates grew significantly faster and to higher total infectious virus titers than A.3. All A.2.5 lineage isolates grew significantly faster at 37°C than at 33°C irrespective of cell type, and to higher peak titers except compared to A.3. This suggests A.2.5’s adapted to improve replication using similar mutations found in the B-lineage SARS-CoV-2 variants.

**Importance:** While both A- and B-lineage SARS-CoV-2 variants emerged and circulated together during the early months of the pandemic, the B-lineages that acquired Spike D614G eventually outcompeted all other variants. We show that the A-lineage variants eventually evolved mutations including Spike D614G and Spike L452R that improved their in vitro replication in human nasal epithelial cells in a temperature dependent manner, suggesting there are some highly selectable mutation landscapes that SARS-CoV-2 can acquire to adapt to replication and transmission in humans.

## Introduction

As of February 2023, the rapid spread of COVID-19 worldwide has caused at least 756 million cases and 6.8 million deaths [1], while driving the emergence of more than 2,000 new PANGO lineages and 30 different Nextstrain clades [2, 3]. The earliest SARS-CoV-2 sequence deposited in GISAID was collected on December 24^th^, 2019 and was designated as a B lineage [4]. Within a week, another isolate was designated as an A lineage with defining amino acid substitutions of L84S in ORF8 and two substitutions C8782T and T28144C [2]. At least two zoonotic events likely caused the split into two root lineages that dominated the first three months of the pandemic: the more ancestrally distant but earlier detected B-lineages (Clade 19A) and the more closely related but later identified A-lineages (Clade 19B) [3, 5, 6].

Despite the higher fidelity of SARS-CoV-2 genome replication due to the presence of the exoribonuclease (nsp14) [7–9], mutations throughout the genome led to the evolution of the B.1-lineages (Clade 20A) and selection of its defining D614G substitution in the Spike gene. More descendent lineages bearing the D614G substitution were rapidly introduced due to the selective advantage conferred by the increased spike stability, viral infectivity, replication, and transmissibility of this substitution [10–13]. The subsequent prominence of these B-lineage variants drove the detection of A-lineage viruses to less than 1% of global sequences by mid-2020 [3].

However from early-2021, the circulation of A-lineages rebounded to 4% of global sequences and was sustained until mid-2021 [3]. The re-emergence of A-lineage variants correlated with the acquisition of new mutations and lineages. A.27 variants were identified in Germany and France, A.23.1 in Uganda and UAE, and A.2.5 in Panama, Canada and the US [14–19]. The A.2.5 variants were also isolated in clinical samples from Johns Hopkins Hospital and Stanford Health Care throughout 2021. This lineage acquired spike mutations including D614G, L452R, Δ141-143, D215A and ins215AGY, some of which are known to confer fitness advantages in other lineages [15, 16, 20]. The D614G substitution is found in all variants of concern (VOCs), while L452R, demonstrated to improve viral infectivity, fusogenicity and evasion of cellular and humoral immunity [21–23], is found in both Delta (B.1.617.2) and Omicron (BA.4, BA.5) variants. Spike 141-144 region is often referred to as the RDR2 (recurrent deletion region 2) in the N-terminal domain (NTD) and deletions in this region are associated with neutralizing escape [24, 25]. The Alpha variant also possesses a similar Δ144 in the RDR2 [26], collectively indicating signs of A.2.5 convergent evolution with VOCs in the spike gene.

This study conducted an *in vitro* characterization of an early A-lineage variant A.3 and three A.2.5 isolates with different mutations to see if and what replication fitness advantages have been conferred by the convergent evolution of A.2.5 variants with B-lineage viruses and VOCs in the spike gene.

## Results

### Continued circulation and acquisition of mutations in A-lineage variants

Corresponding to the timeframe of A-lineage global rebounds, A.2.5 lineage variants and an A.23.1 variant was detected in early to mid-2021 at Johns Hopkins Hospital **(Fig. 1)**. Early A-lineage variants, such as A, A.1, A.2.2 and A.3, were readily detected up to mid-2020, accounting for 20% of Johns Hopkins Hospital COVID-19 cases in March 2020. No A-lineages were detected until when A.2.5 variants emerged in 2021. An A.3 strain (HP00076) and three A.2.5 strains (HP02153, HP05922 and VRLC091) were isolated **(Table 1)** and whole genome sequencing revealed the A.2.5 viruses accumulated mutations throughout the genome when compared to A.3, with each isolate encoding a unique mutation profile **(Fig. 2).** At least five conserved spike mutations were seen across the A.2.5 isolates: Δ141-143, D215A, ins215AGY, L452R and D614G. VRLC091 uniquely had acquired three additional substitutions W152R, T240I and A263S. These mutations, found similarly in other VOCs **(Table 2)**, represent signs of convergent evolution of A.2.5 variants with the spike sequences of VOCs.

**Figure 1.**
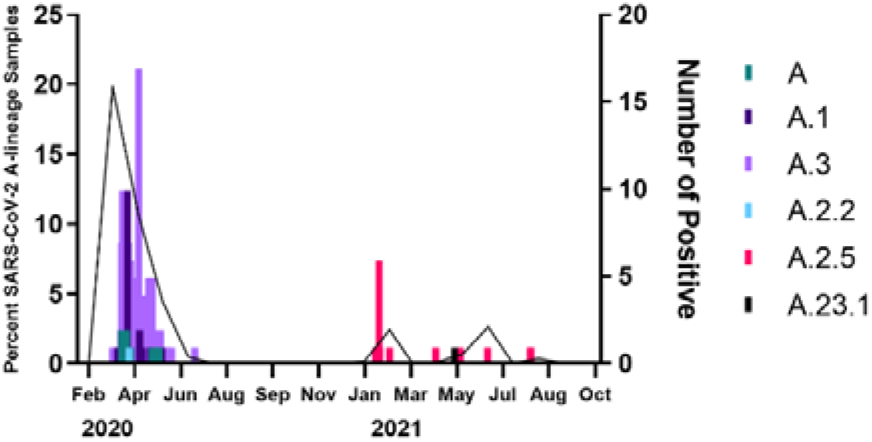
Detection of A-lineage (Clade 19B) SARS-CoV-2 variants from the Johns Hopkins Hospital. 209 A-lineage (Clade 19B) isolates were characterized from the Johns Hopkins Health System samples from March 2020 to August 2021. The total A-lineage (Clade 19B) percent positivity per month is represented in lines over time (left Y-axis), while the number of positive cases for each A-lineages are presented in bars (right Y axis)

**Table 1.** A-lineage clinical isolates from Johns Hopkins Health System and Stanford Health Care.

| Lineage | Strain | GISAID Accession Number | Date Collected | Institution Collected | Patient Information (Sex/Age) |
| --- | --- | --- | --- | --- | --- |
| **A.3** | hCoV-19/USA/MD-HP00076/2020 | EPI_ISL_438234 | 2020-03-11 | Johns Hopkins Hospital | Male/30 |
| **A.2.5** | hCoV-19/USA/MD-HP02153/2021 | EPI_ISL_981090 | 2021-02-02 | Johns Hopkins Hospital | Female/18 |
| **A.2.5** | hCoV-19/USA/MD-HP05922/2021 | EPI_ISL_2331556 | 2021-05-09 | Johns Hopkins Hospital | Female/38 |
| **A.2.5** | hCoV-19/USA/CA-VRLC091/2021*** | EPI_ISL_3050209 | 2021-06-29 | Stanford Health Care | Female/23 |

\*\*\*Also named as hCoV-19/USA/CA-Stanford-31\_S39/2021 on GISAID database.

**Figure 2.**
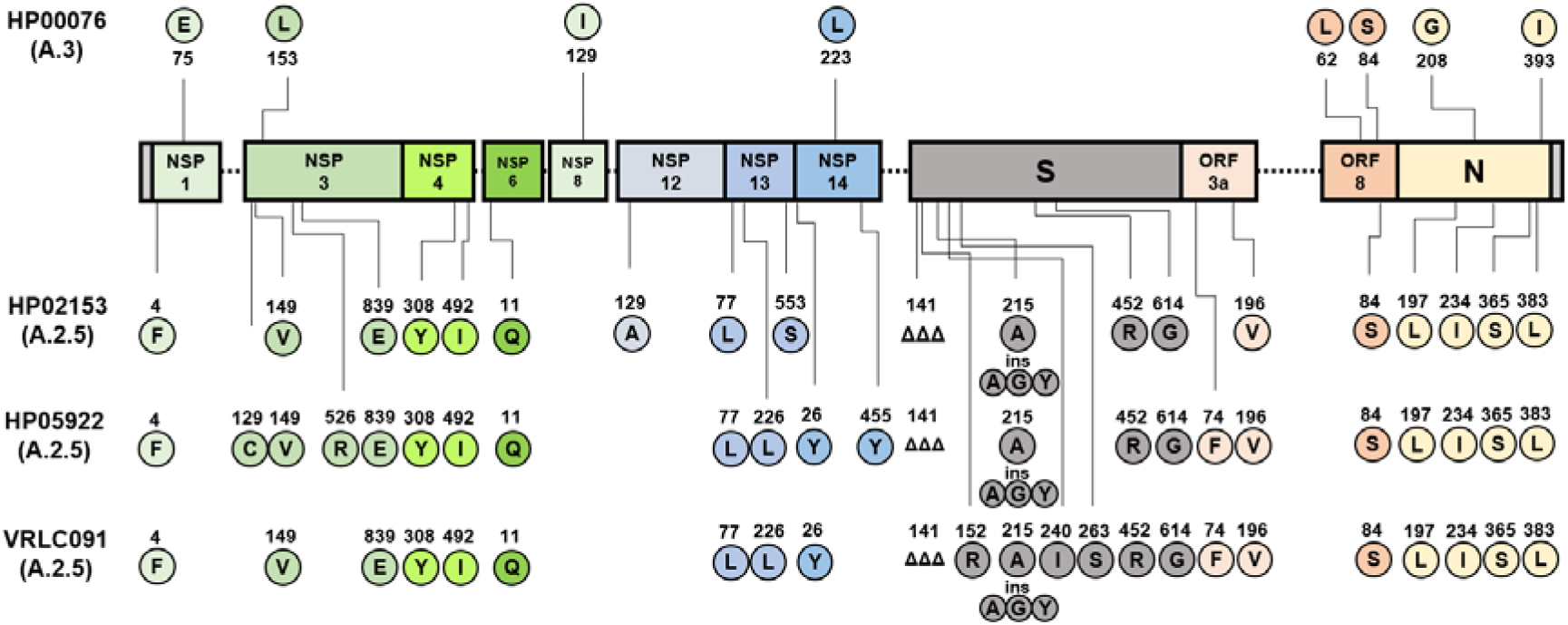
Mutation comparison of A-lineage clinical isolates. A-lineage isolate-specific mutations were compared to Wuhan/Hu-1 as reference strain.

**Table 2.** A.2.5 isolates acquired similar or same spike mutations as VOCs.

| PANGO Lineage | Virus Name | Accession # | Spike (S1) |  |  |  |  |  |  |
| --- | --- | --- | --- | --- | --- | --- | --- | --- | --- |
| B | Wuhan/Hu-1/2019 (reference) | EPI_ISL_402125 | NTD |  |  |  |  | RBD | CTD |
|  |  |  | L141 | W152 | D215 | T240 | A263 | L452 | D614 |
| A.3 | HP00076 | EPI_ISL_438234 |  |  |  |  |  |  |  |
| A.2.5 | HP02153 | EPI_ISL_981090 | ΔΔΔ |  | A,AGY ins |  |  | R | G |
| A.2.5 | HP05922 | EPI_ISL_2331556 | ΔΔΔ |  | A,AGY ins |  |  | R | G |
| A.2.5 | VRLC091 | EPI_ISL_3050209 | ΔΔΔ | R | A,AGY ins | I | S | R | G |
| B.1.1.7 | Alpha | EPI_ISL_825013 | Y144Δ |  |  |  |  |  | G |
| B.1.351 | Beta | EPI_ISL_890360 |  |  | G | L241 ΔΔΔ |  |  | G |
| AY.106 (B.1.617.2) | Delta | EPI_ISL_2331507 | G142D | E156G,ΔΔ | G |  |  | R | G |
| P.1 | Gamma | EPI_ISL_2331506 | D138Y |  |  |  |  |  | G |
| BA.1.18 (B.1.1.529) | Omicron | EPI_ISL_7160424 | G142 D,ΔΔΔ |  | 214 EPE ins |  | D |  | G |
| BA.4 (B.1.1.529) | Omicron | EPI_ISL_12416220 | G142D |  | V213G |  | D | R | G |

### A.2.5 escapes neutralization from COVID-19 convalescent plasma compared to A.3

The early A.3 lineage isolate and later A.2.5 lineage isolates were compared for their ability to escape neutralizing antibody responses. Plaque reduction neutralization tests (PRNT) against the four isolates were conducted with a panel of ten COVID-19 convalescent plasma (CCP) samples collected from May 2020 and March 2021 (between A.3 and A.2.5 collection). All three A.2.5 lineage viruses showed enhanced escape from neutralization when compared to the A.3 isolates **(Fig. 3)** with a reduction in geometric mean IC_50_ titer (GMT) ranging from 2.5-to 5-fold when compared to the GMT of A.3. This data indicates that A.2.5 isolates had evolved to partially escape neutralizing activity generated against viruses circulating during the same timeframe.

**Figure 3.**
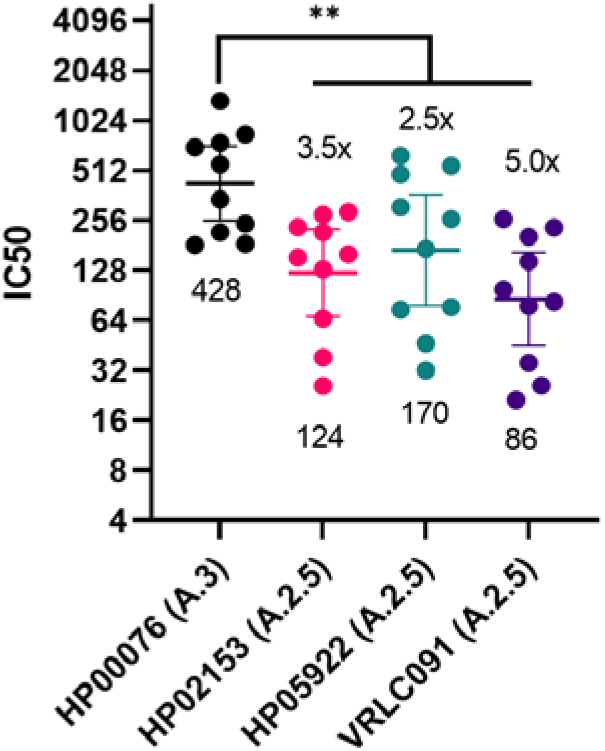
Plaque reduction neutralization test (PRNT) against A-lineage isolates. Ten convalescent plasma samples from patients previously infected with SARS-CoV-2 between May 2020 and March 2021 were tested against each of the A-lineage isolates to determine virus neutralization activity in VeroE6-TMPRSS2 cells. IC_50_ (half maximal inhibitory concentration) was calculated using the inhibition-dose response of plaque reduction over 2-fold serial dilutions of plasma starting from 1:20. Geometric mean titer and 95% confidence intervals are shown per isolate, and the geometric mean fold drops of A.2.5 isolates relative to that of HP00076 (A.3) is denoted above each data set and the geometric mean titer indicated below. Statistical significance was measured using the Wilcoxon matched-pairs signed rank test, ** p=0.002.

### A.2.5 generate smaller plaque sizes than A.3 at both 33°C and 37°C on VeroE6 cells

Plaque assays were conducted on VeroE6 cells at both 33°C and 37°C. By 96hpi at 37°C and by 120hpi at 33°C, all four isolates generated countable and opaque plaques (**Fig. 4A-B**). Measuring and comparing the plaque areas at 37°C revealed that all three A.2.5 isolates generated smaller mean (±SD) plaque sizes (HP02153, 2.50±0.90mm^2^; HP05922, 3.26±1.42mm^2^; VRLC091, 4.28±1.67mm^2^) than the A.3 isolate HP00076 (6.42±2.45mm^2^; **Fig. 4C**). A.2.5 isolates generated mean plaque areas that were 1.5 to 2.6-fold smaller than A.3, while VLRC091 formed the largest plaques among the A.2.5 isolates. At 33°C, all A.2.5 isolates (HP02153, 2.28±0.76mm^2^; HP05922, 3.24±0.88mm^2^; VRLC091, 3.70±1.69mm^2^) also produced significantly smaller plaques than HP00076 (8.01±2.87mm^2^), with 2.2 to 3.5-fold smaller plaques than A.3 (**Fig. 4D**).

**Figure 4.**
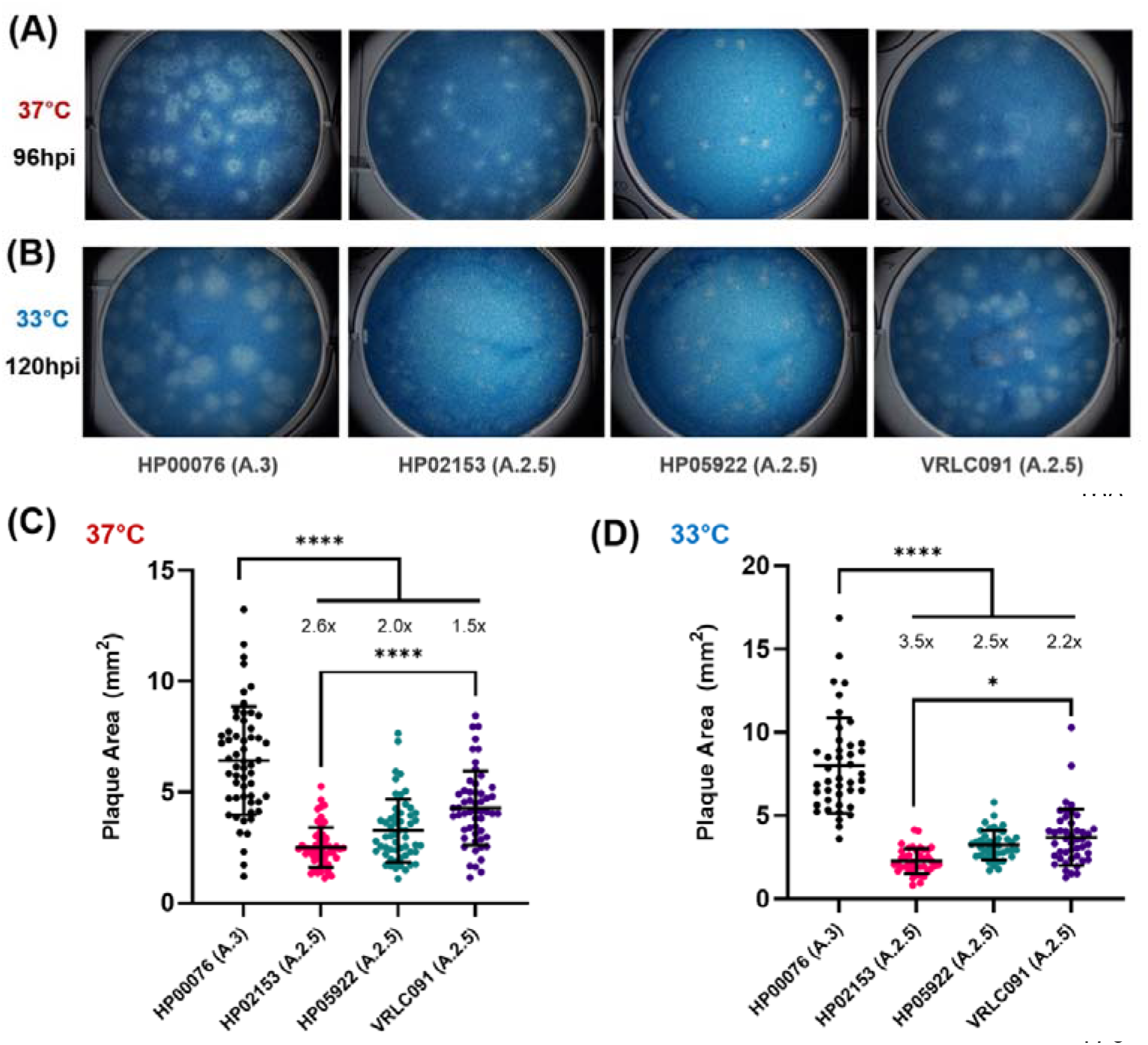
Methylcellulose plaque assay of A-lineage isolates grown on VeroE6 cells at 33°C and 37°C. Methylcellulose overlay plaque assays were conducted on VeroE6 monolayers with A-lineage isolates at both 33°C and 37°C. **(A)** Representative plaque images of A-lineage isolates at 37°C and **(B)** at 33°C. Infections were incubated at 33°C for 120hpi and at 37°C for 96hpi for most distinguishable plaques. **(C)** A-lineage isolate plaque areas at 37°C and **(D)** at 33°C. Data was generated from at least 40 countable plaques across three independent experiments for each virus and temperature, and areas quantified on ImageJ. Ordinary one-way ANOVA with Tukey’s multiple comparison test was used to determine statistical significance, **** p<0.0001, * p=0.0240. Mean plaque area fold decreases were calculated for each virus relative to the mean plaque area of HP00076.

### A.2.5 generates smaller plaques than A.3 on VeroE6-TMPRSS2 overexpressed cells at 33°C and 37°C

To determine the effects of increased expression of TMPRSS2 (transmembrane serine protease 2) – a protease implicated in the processing and fusogenicity of SARS-CoV-2 Spike protein - a plaque assay was also conducted on VeroE6-TMPRSS2-overexpressed cell monolayers. In contrast to the opaque plaques seen on VeroE6 cells, all the isolates generated clear and easily distinguishable plaques by 48hpi at 37°C and by 72hpi at 33°C (**Fig. 5A-B**). While observable plaques formed more quickly in VeroE6-TMPRSS2 cells than on WT VeroE6 cells, A.2.5 isolates still formed smaller plaques than the A.3 isolate at both temperatures (**Fig. 5C-D)**. At 37°C, A.2.5 isolates’ plaques (HP02153, 0.74±0.27mm^2^; HP05922, 0.67±0.34mm^2^; VRLC091, 0.65±0.35mm^2^) were 2.6-3.0-fold smaller than those of A.3 (1.94±1.00 mm^2^; **Fig. 5C**). At 33°C, A.2.5 isolates’ plaques (HP02153, 0.27±0.12mm^2^; HP05922, 0.31±0.19mm^2^; VRLC091, 0.53±0.37mm^2^) were 2.3-4.4-fold smaller than those of A.3 (1.21±0.60mm^2^; **Fig. 5D**). Taken together, the A.2.5 isolates showed smaller plaques when compared to an earlier A.3 isolate at both 33°C and 37°C, and in VeroE6 cells that expressed low or high amounts of TMPRSS2.

**Figure 5.**
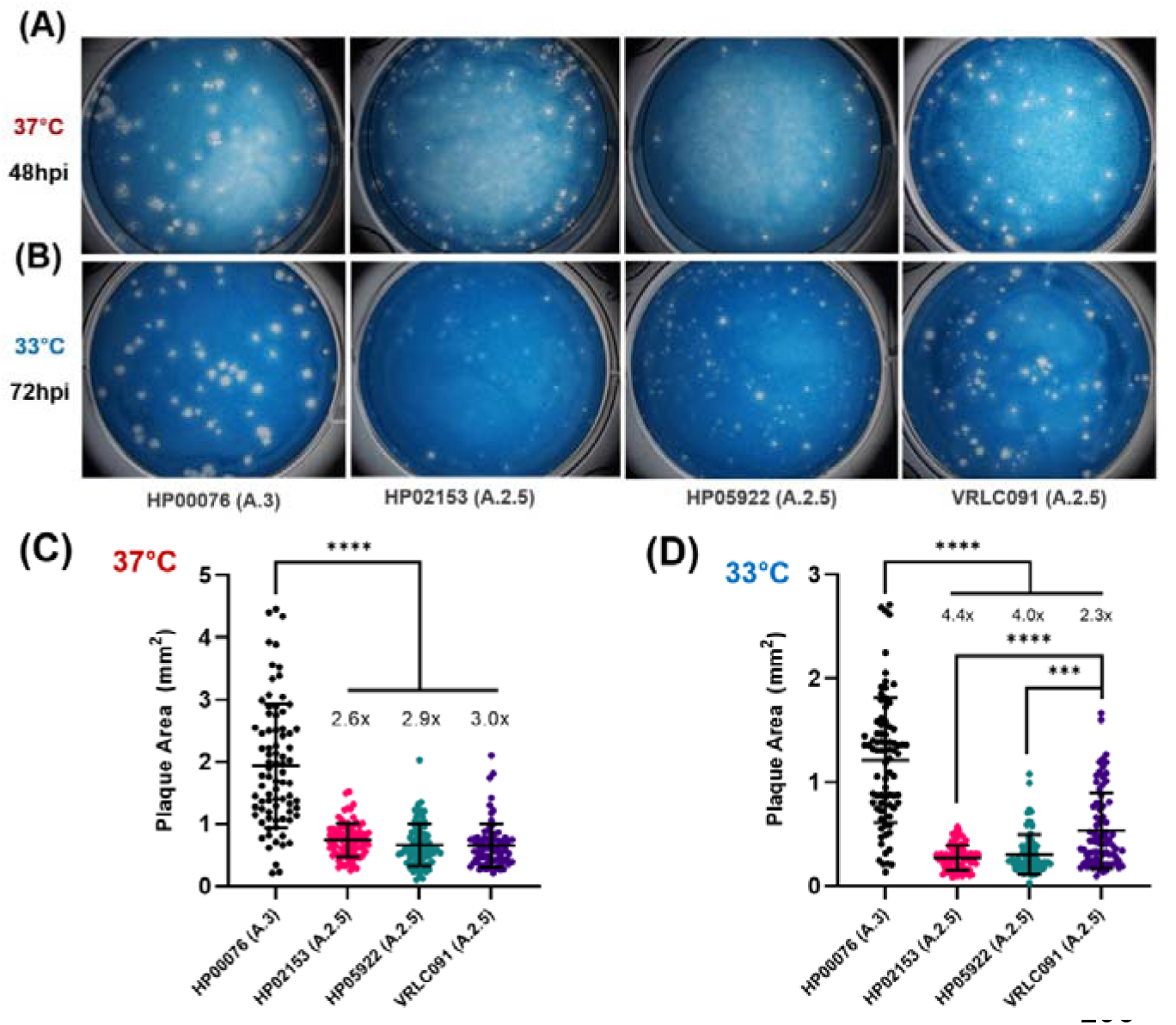
Methylcellulose plaque assay of A-lineage isolates on VeroE6-TMPRSS2 cells at 33°C and 37°C. Methylcellulose overlay plaque assays were conducted on VeroE6-TMPRSS2 overexpressed cell monolayers with A-lineage isolates at both 33°C and 37°C. **(A)** Representative plaque images of A-lineage isolates at 37°C and **(B)** at 33°C. Infections were incubated for 48hpi at 37°C and for 72hpi at 33°C. **(C)** A-lineage isolate plaque areas at 37°C and **(D)** at 33°C. Data for 37°C and 33°C each were accumulated from 80 observable plaques measured from three independent experiments for each virus and temperature, and areas measured using ImageJ. Statistical significance was measured using ordinary one-way ANOVA with Tukey’s multiple comparison test, **** p<0.0001, *** p=0.0006. The mean plaque area fold drops were calculated for each virus relative to the mean plaque area of HP00076 and denoted above data points.

### A.2.5 grows slower and produces lower total infectious viruses than A.3 on VeroE6-TMPRSS2 cells

To further characterize virus replication, VeroE6-TMPRSS2 cells were infected with A-lineage isolates at a low MOI of 0.01 TCID_50_/cell at 33°C and 37°C. At 33°C, HP00076 (A.3) infectious virus production was significantly faster than the A.2.5 isolates with a steeper slope and faster time to peak virus titer of 72hpi, compared to HP02153 (A.2.5), HP05922 (A.2.5), and VRLC091 (A.2.5) which achieved peak titer at 96hpi or after (**Fig. 6A**). VRLC091 grew fastest among the A.2.5 isolates, but slower than HP00076. At 37°C, the A.3 isolate infectious virus production was also faster than what was observed with HP02153 and HP05922 but was not different from VRLC091 **(Fig. 6B)**. These replication differences were somewhat consistent with the plaque assay data, where the A.3 isolate formed larger plaques than all of the A.2.5 isolates, and VRLC091 showed slightly larger plaques compared with the other A.2.5 isolates at 33°C **(Fig. 5D)**. Differences in growth curves were less apparent at 37°C compared to 33°C, and also indistinguishable when comparing the total area under the curve (AUC) calculated from each replicate well, which represents the total amount of infectious virus produced across the growth curve (**Fig. 6C)**. At 33°C, the A.3 isolate still produced significantly more total infectious viruses than the A.2.5 isolates, with VRLC091 the most among the A.2.5 isolates **(Fig. 6C)**. When the growth of each isolate was compared across temperatures, all isolates grew significantly faster at 37°C than at 33°C by reaching faster peak titers **(Fig. 6D)**.

**Figure 6.**
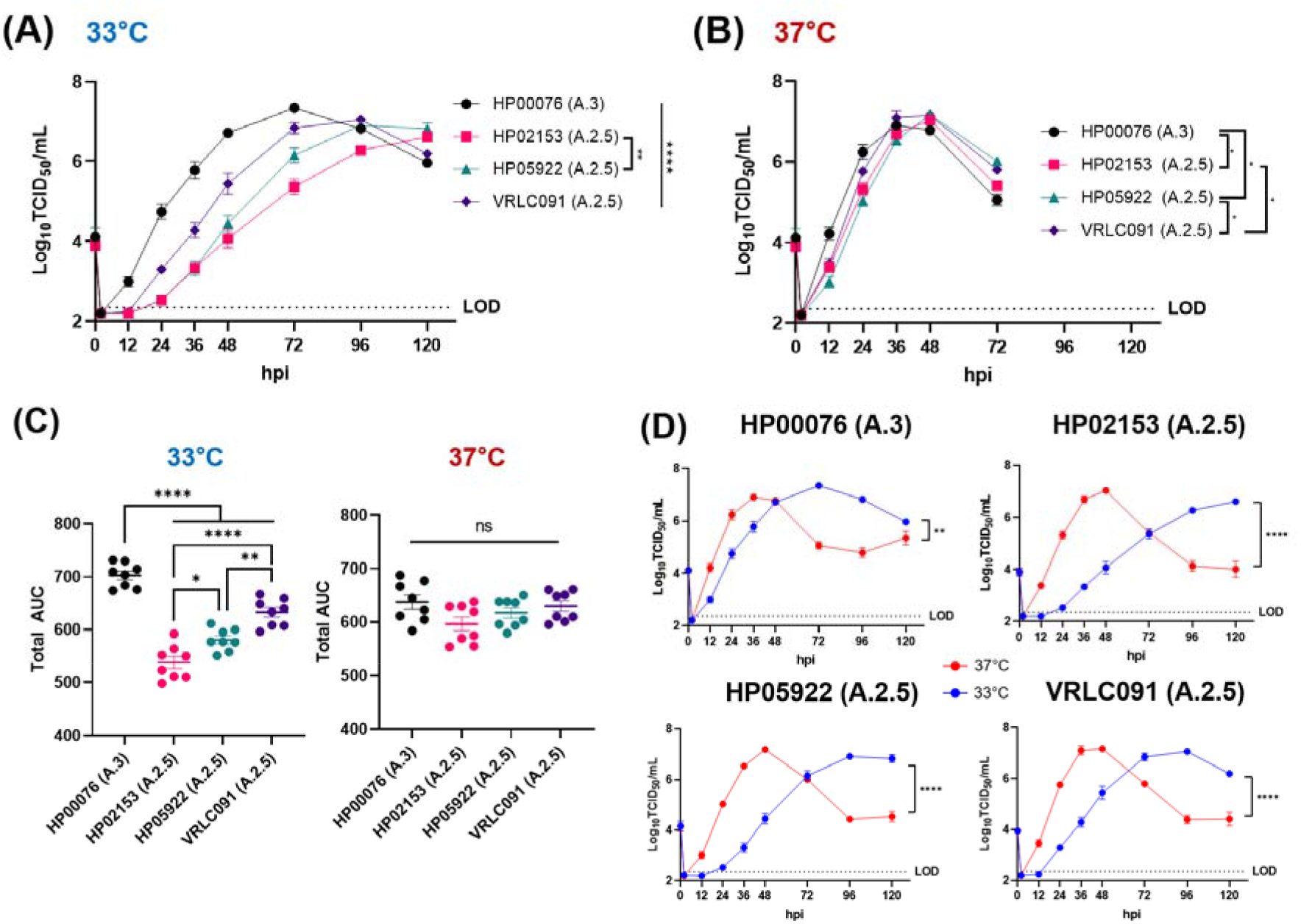
Temperature-dependent low MOI growth curve infections with A-lineage isolates on VeroE6-TMPRSS2 cells. VeroE6-TMPRSS2 cells were infected at an MOI of 0.01 with A-lineage isolates at both 33°C and 37°C. **(A)** VeroE6-TMPRSS2 growth curve infection at 33°C. **** p<0.0001, ** p=0.0083. **(B)** VeroE6-TMPRSS2 growth curve infection at 37°C. * p=0.0436 (HP00076vsHP02153), p=0.0245 (HP00076vsHP05922), p=0.0245 (HP02153vsVRLC091), p=0.0132 (HP05922vsVRLC091). Titers from two independent growth curve infections with four replicate wells per virus at each temperature were collapsed for the data. LOD (limit of detection) of TCID_50_ assay = 2.37 logTCID_50_/ml. Statistical significance was calculated using two-way ANOVA with Tukey’s multiple comparison test. **(C)** Total AUC of replicate wells from 33°C and 37°C VeroE6-TMPRSS2 growth curves. Total AUC was calculated for each replicate well from the 33°C and 37°C growth curves in (A) and (B) with the standard error of mean (SEM) using GraphPad Prism 9 software. One-way ANOVA with Tukey’s multiple comparison test was used to determine statistical significance, * p=0.0118, ** p=0.0022, **** p<0.0001. **(D)** VeroE6-TMPRSS2 growth curve comparisons by temperature. 33°C is denoted in blue and 37°C is represented in red. Two-way ANOVA with Tukey’s multiple comparison test was conducted for statistical significance, ** p=0.0057, **** p<0.0001.

### A.2.5 isolates undergo more syncytia formation than A.3 but possess varying spike cleavage efficiencies in VeroE6-TMPRSS2 cells

Images of representative wells from the VeroE6-TMPRSS2 growth curves (**Fig. 6**) were also taken to monitor for cytopathic effect (CPE) differences during infection. A.2.5 isolate infections displayed significantly more cell clumping compared to A.3 at both 33°C and 37°C (**Fig. 7A-B**). A.2.5 isolates formed visibly larger and darker aggregates of cells and, at a higher total magnification of 200X, HP00076 exhibited minimal cell clumping, while all the A.2.5 isolates formed large clumps of cells (**Fig. 7C**).

**Figure 7.**
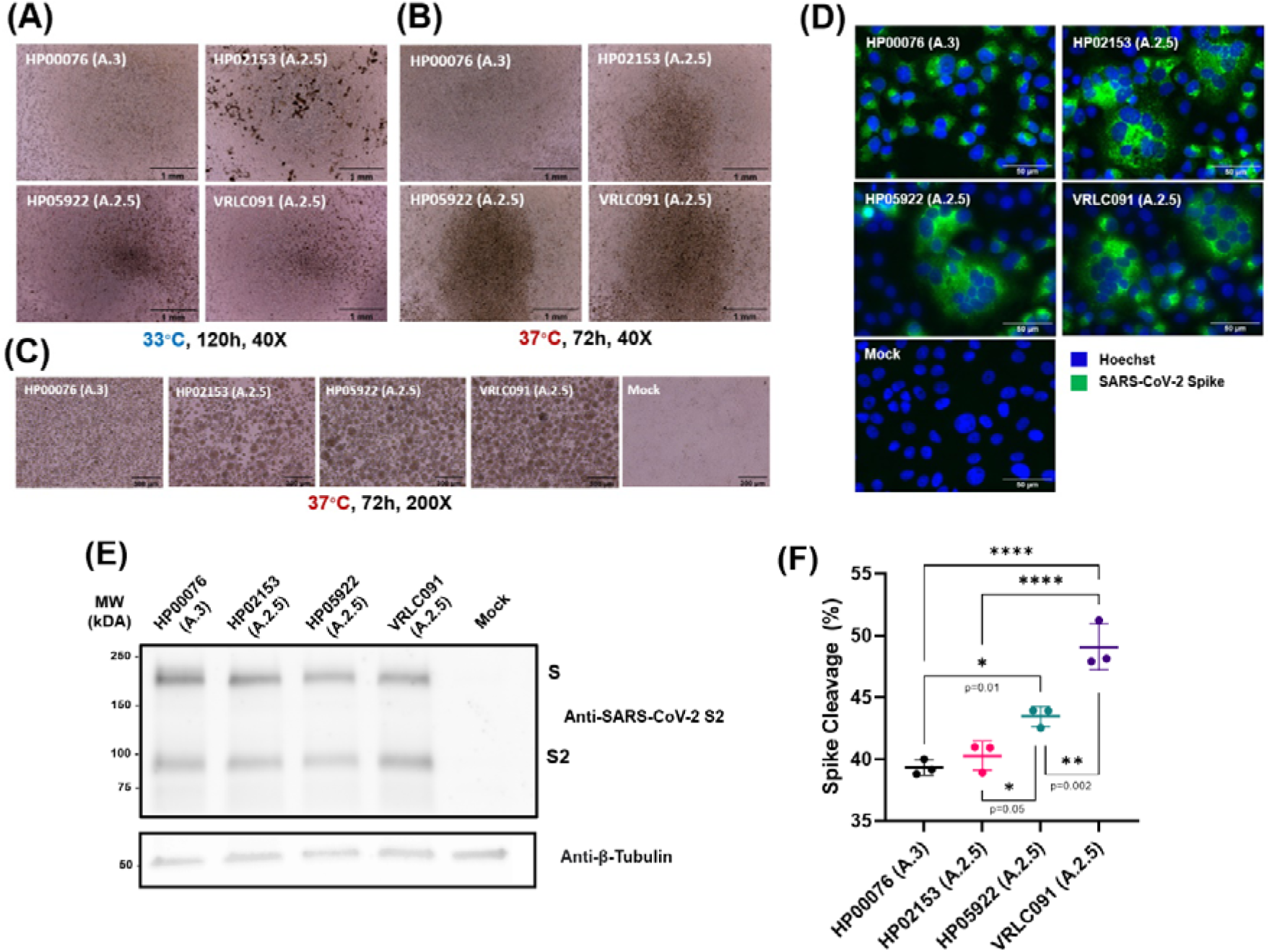
Fusion capabilities of A-lineage isolates represented through CPE images, immunofluorescence imaging and spike cleavage. Representative wells were imaged for CPE over the course of VeroE6-TMPRSS2 infections described in Figure 6 using a brightfield microscope. **(A)** CPE images from representative wells at 33°C after peak CPE detection. A.2.5 isolates showed maximal CPE at 120hpi, while HP00076 exhibited peak CPE at 96hpi. Images were taken at 40X total magnification. Scale bar, 1mm. **(B)** CPE images from representative wells at 37°C after peak CPE detection. A.2.5 isolates produced the most CPE at 72hpi while HP00076 demonstrated peak CPE at 48hpi. Images were also taken at 40X total magnification. Scale bar, 1mm. **(C)** Representative CPE images at 200X total magnification. These images are 200X magnifications of the same four infected wells from (D) with an uninfected mock. Scale bar, 300μm. VeroE6-TMPRSS2 cells were also infected at a higher MOI of 1 with A-lineage isolates for immunofluorescence imaging and for western blotting of the spike protein to determine spike cleavage levels. **(D)** Immunofluorescence confocal images of VeroE6-TMPRSS2 cells infected with A-lineage isolates at 37°C. VeroE6-TMPRSS2 cells were stained using anti-Spike protein antibody (green) and Hoechst (blue). Representative images were taken using a Widefield microscope at 400x total magnification. Scale bar, 50μm. **(E)** Immunoblotting of VeroE6-TMPRSS2 cell lysates infected with A-lineage isolates for spike cleavage. Spike expression levels were determined with anti-SARS-CoV-2 S2 antibodies. Representative gel of one replicate is shown. Under each lane of the blot are the average spike cleavage percentage and standard error of mean (SEM) produced by three independent replicates. Anti-beta-tubulin staining is also shown for loading control. **(F)** Spike cleavage efficiency per A-lineage isolates. Spike cleavage efficiency calculated from three replicate western blots of cell lysates are plotted for average spike cleavage percentage and SEM, as indicated in (E). Statistical analyses were conducted using ordinary one-way ANOVA with Tukey’s multiple comparison test. Aside from **** p<0.0001, p-values are indicated below each asterisk

Cell clumping could indicate either syncytia formation or aggregation of cells by transitory cell-to-cell interactions. To more carefully differentiate between these two outcomes, immunofluorescence imaging was performed on cells infected with each of the A-lineage isolates to distinguish syncytia. Images revealed that A.2.5 isolates all formed substantial syncytia as indicated by SARS-CoV-2 spike positive and multinucleated cells (**Fig. 7D**). HP00076 (A.3) did not form significant syncytia.

To determine if Spike cleavage efficiency was driving the difference between the A.3 and A.2.5 isolates, VeroE6-TMPRSS2-infected cells were harvested for Western blotting. Immunoblotting for the S2 domain of the spike protein generated two bands per lane in infected samples representing the full-length spike (S) and the S2 domain (**Fig. 7E**). Analysis of the S and S2 protein expression levels suggested that HP05922 and VRLC091 achieved higher mean spike cleavage efficiency of 43.46±0.36% (±SEM) and 49.09±1.08% compared to HP00076’s significantly lower 39.32±0.36% (**Fig. 7F**). HP02153 was only marginally higher at 40.29±0.68%. The variability in Spike cleavage efficiency, particularly of HP02153, suggests this alone was not driving the differences in syncytia formation seen between A.3 and A.2.5 isolates.

### A.2.5 isolates replicate faster and produce more infectious virus than A.3 on hNEC cultures

A-lineage isolate replication was also assessed on human nasal epithelial cell (hNEC) cultures to determine viral growth kinetics on a more physiologically relevant cell culture model. The hNEC cultures were infected at an MOI of 0.1 and infections performed at both 33°C and 37°C. At 33°C, minimal growth differences were observed between A.2.5 and A.3 with only HP05922 showing faster kinetics of infectious virus production compared to HP00076 (**Fig. 8A**). When the total AUC of each of the replicate wells were calculated to determine total infectious virus production, no statistical differences were seen between the isolates (**Fig. 8C**). However, at 37°C, a clear growth difference was seen with A.2.5 isolates growing more rapidly than A.3 and achieving almost ten-fold (1 log_10_) higher peak titers albeit at approximately the same 96hpi peak time for all isolates (**Fig. 8B**). HP00076 reached a lower mean peak-titer (±SEM) of 6.794±0.244 logTCID_50_/ml compared to HP02153’s 7.678±0.128 logTCID_50_/ml, HP05922’s 8.074±0.121 logTCID_50_/ml, and VRLC091’s 8.241±0.199 logTCID50/ml. All A.2.5 isolates produced higher amounts of total infectious virus than A.3 with HP00076 having a total AUC (±SEM) of 905.2±20.48, compared to the A.2.5 isolates HP02153 (1063±7.101), HP05922 (1049±9.116) and VRLC091 (1060±12.73; **Fig. 8C**). Furthermore, when growth curves were stratified for temperature, all A-lineage isolates grew more rapidly at 37°C than at 33°C. Generally, all isolates reached peak titers at around 96hpi when incubated at 37°C, while peak titers at 33°C were achieved by 144hpi or beyond (**Fig. 8D**). Although growing slower at 33°C, A.3 isolate HP00076 still reached higher peak titer at 33°C than at 37°C while A.2.5 isolates generated faster and higher peak titers at 37°C (**Fig. 8D**). Taken together, the A.2.5 isolates showed equivalent or better infectious virus production compared to the A.3 lineage in hNEC cultures, indicating that the accumulation of mutations in A.2.5 isolates results in viruses with improved replication fitness, particularly at 37°C in hNEC cultures.

**Figure 8.**
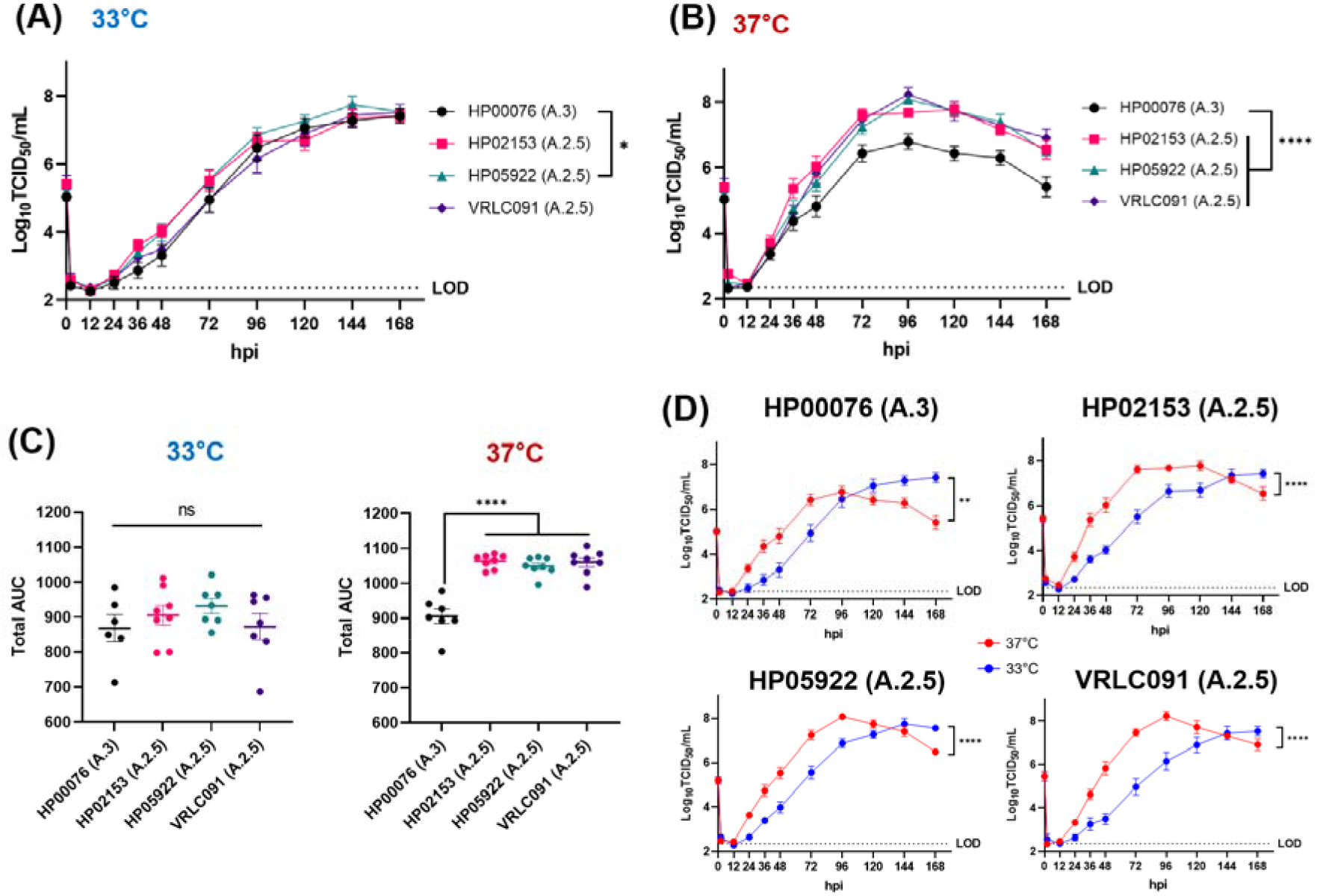
Low MOI hNEC growth curve infections of A-lineage isolates at 33°C and 37°C. hNECs were infected with A-lineage isolates at 0.1 MOI to observe growth trends at 33°C and 37°C. **(A)** hNEC growth curve infection at 33°C, * p=0.0278. **(B)** hNEC growth curve infection at 37°C, **** P<0.0001. Data for each temperature were accumulated from two independent experiments with four replicate wells per virus. Statistical significance was determined using two-way ANOVA with Tukey’s multiple comparison test. LOD (limit of detection) of TCID_50_ assay = 2.37 logTCID_50_/ml. **(C)** Total AUC of all replicate wells from 33°C and 37°C hNEC growth curves. Total AUC was calculated for each replicate wells from the 33°C and 37°C growth curves in (A) and (B) with the standard error of mean (SEM) using GraphPad Prism 9 software. One-way ANOVA with Tukey’s multiple comparison test was used to determine statistical significance for differences, **** p<0.0001. **(D)** hNEC growth curve comparisons by temperature. Data from (A) and (B) were combined to generate temperature-dependent comparisons. Two-way ANOVA with Tukey’s multiple comparison test was conducted for statistical significance, ** p=0.0037, **** p<0.0001.

## Discussion

Convergent evolution of the spike gene has been an ongoing phenomenon and is especially seen among VOCs. The independent emergence of Alpha, Beta and Gamma variants in late 2020 to early 2021 demonstrated convergent evolution of the Spike gene and positive selection for mutations like N501Y and E484K [27], and later in Delta and among Omicron sub-lineages such as with ΔHV69-70, L452R, E484K and P681H mutations [28, 29]. Convergent evolution is also seen with A.2.5 variants which is believed to have evolved independently but did not necessarily reach the same global circulation as did by VOCs that are descendants of the B-lineages. Although A.2.5 variants never reached global prominence, convergent evolution of the Spike gene had likely provided sufficient fitness advantage to allow their rebound in early 2021 and compete amidst the emergence and global presence of VOCs such as Alpha and Delta variants.

Live virus, *in vitro* characterization experiments compared three A.2.5 lineage isolates, collected during the timeframe corresponding to the re-emergence of A-lineage SARS-CoV-2 variants, with an early A.3 isolate collected in March 2020. PRNTs indicated the A.2.5 variants acquired neutralization escape capabilities compared to the more earlier A.3 strain, which is in line with reported reduced plasma recognition by A.2.5 Spike and neutralization capacity compared to the ancestral D614G strain [30]. Infections of more physiologically relevant model of hNEC cultures also demonstrated that the A.2.5 variants produced 10-fold more total viruses during the course of infection and replicated at a faster rate than A.3 at 37°C, which may contribute to more viral load in patients. This observation could further be supported as the D614G Spike substitution, present in A.2.5 but not in A.3, has been found to restore trafficking of Spike to lysosomes and enhance the use of the cathepsin L pathway to increase SARS-CoV-2 infectivity and stability at 37°C [31].

However, this growth trend observed in hNEC cultures was not observed in VeroE6 cell lines in both WT and TMPRSS2-overexpressed cell lines and was in fact the opposite. This implies that newly acquired A.2.5 mutations may be playing a key role in mediating fitness advantages in primary nasal epithelial cell cultures, but not in immortalized cells like VeroE6 which also lack interferon responses [32] and may have permitted unrestricted growth even to early variants like A.3. It is unclear what other specific mutations are driving the improved replication of A.2.5 isolates in hNEC cultures, and the contribution of both Spike and non-Spike mutations needs to be investigated more thoroughly. Nonetheless, this finding also highlights the importance of appropriate cell culture model selection to characterize viral growth kinetics, as the properties of immortalized cell lines do not fully mimic the physiological conditions.

In VeroE6-TMPRSS2 cells, the slower spreading and growing A.2.5 isolates also correlated to significant syncytia formation, while the A.3 isolate, which grew faster, failed to produce syncytia. Taken together with the low MOI growth curve data, it is possible that increased syncytia formation is potentially delaying release or spread of progeny viruses, thereby preventing subsequent infection of neighboring cells and reducing overall infectious virus production. Spike mutations were likely driving differences in syncytia formation, however it was not clearly mediated by improved spike cleavage efficiency as seen with variable increases in efficiency among A.2.5 isolates compared to that of A.3. Other factors like differences in Spike binding affinity for ACE2 and increased cell surface levels of Spike protein might also explain the enhanced syncytia formation seen in A.2.5 isolates.

Another significant observation was that growth-related experiments indicated that all A-lineage isolates, irrespective of their presence or absence of Spike mutations, in both VeroE6 cell lines and in primary hNEC cultures grew faster at 37°C. A.2.5 isolates particularly benefited growth at 37°C as they propagated faster and to higher titers than at 33°C in hNECs. The early A.3 isolate also grew faster at 37°C but did not seem to reach higher peak titers than at 33°C in hNECs, making the earlier strain less obvious in terms of temperature preference. Overall, this contrasts with other reported data that indicate faster and higher *in-vitro* SARS-CoV-2 replication efficiency at 33°C and SARS-CoV replication at 37°C in human tracheobronchial epithelial cells [33]. Elevated respiratory tract temperatures are known to impact and restrict replication of respiratory viruses, such as SARS-CoV-2, by optimal induction of interferon and interferon-independent antiviral immune defenses [33, 34], and negatively impact ACE2 binding by SARS-CoV-2 Spike [30, 35]. The A-lineage variants, phylogenetically closer to ancestral bat sarbecoviruses [5], may possess key mutations in the non-structural genes not present in B-lineage variants that dictate replication tolerance and efficiency for core body temperatures such as the lower respiratory tract, which are characteristic of spillover viruses with lower transmission and more disease severity [36].

Spike mutations are likely contributing to differences in virus properties such as growth in primary hNEC cultures, neutralization escape and in syncytia formation, and infectious clone technologies could be used to identify mutations that drive these differences. Nevertheless, the findings of A.2.5 neutralization escape to circulating, pre-existing immunity and overall enhanced replication capabilities in hNEC cultures support the re-emergence of A-lineage variants. Furthermore, temperature adaptation of A-lineage variants, especially A.2.5, to 37°C, which is representative of the lower respiratory tract temperature or warmer upper respiratory tract temperatures during fevers, could have improved viral fitness especially in the context of human pathogenesis. However other factors such as transmission efficiency, competition with fitter variants, and lack of sustained transmission may have also played important roles in explaining the eventual disappearance of A-lineage variants from circulation in humans even before the emergence of the Omicron variants.

## Methods

### Biosafety Containment

All experiments with SARS-CoV-2 viruses were performed under institution-approved biosafety protocols in Biosafety Level 3 (BSL-3) containment.

### Institutional Review Board Approvals

For convalescent plasma, clinical specimen were obtained with written informed consent per the protocols approved by the institutional review boards at Johns Hopkins University School of Medicine as single Institutional Review Board for all participating sites and the Department of Defense Human Research Protection Office. For virus isolation, nasal swabs from SARS-CoV-2 infected individuals were obtained and sequenced with a waiver of consent under the Johns Hopkins protocol number IRB00221396. Virus isolation was performed on deidentified samples under Johns Hopkins protocol number IRB00288258.

### Cell Lines

VeroE6 and VeroE6-Transmembrane Serine Protease 2 (TMPRSS2) overexpressing cells [37] were cultured at 37°C and 5% CO_2_ in complete cell culture media (CM; DMEM supplemented with 1% GlutaMAX (Life Technologies, Cat#35050061), 10% Fetal Bovine Serum (FBS; Gibco, Cat#26140079), 1% Penicillin/Streptomycin mixture (Quality Biologicals, Cat#120-095-721), and 1% 100mM sodium pyruvate solution (Sigma, Cat#S8636-100ML)).

Human nasal epithelial cells (HNEpC; PromoCell, Cat#C-12620) were expanded to confluency with PneumaCult™ Ex Plus Media (StemCell, Cat#05040) at 37°C and 5% CO_2_. Confluent cells were fully differentiated in ALI (air-liquid interface) with PneumaCult ALI Basal Medium (Stemcell, Cat#05002) and 1X PneumaCult ALI Supplement (Stemcell, Cat#05003). 1% PneumaCult ALI Maintenance Supplement (Stemcell, Cat#05006), 0.5% Hydrocortisone stock solution (Stemcell, Cat#07925) and 0.2% Heparin solution (Stemcell, Cat#07980) were added to the ALI Basal Medium.

### Virus Isolation and Virus Seed Stocks

From human nasal swabs collected at Johns Hopkins Hospital, SARS-CoV-2 isolates hCoV-19/USA/MD-HP00076/2020 (GISAID accession: EPI_ISL_438234), hCoV-19/USA/MD-HP02153/2021 (GISAID accession: EPI_ISL_981090) and hCoV-19/USA/MD-HP05922/2021 (GISAID accession: EPI_ISL_2331556) were isolated, while hCoV-19/USA/CA-VRLC091/2021 (GISAID accession: EPI_ISL_3050209) was isolated from a nasal swab from Stanford Health Care [38, 39]. VeroE6-TMPRSS2 cells were grown to 75% confluency in 6-well plates. CM was replaced with 350μL of IM (infection media; same as CM except supplemented with 2.5% FBS). After, 150 μL of the clinical specimen was added to each well and incubated at 37°C for 2 hours. The inoculum was aspirated and replaced with 0.5 mL of IM and cultured at 37°C. The cells were monitored daily for CPE. When CPE was visible in more than 75% of the cells, the IM supernatant was harvested, aliquoted and infectious virus titer determined by TCID50. Virus genome sequence was confirmed to be at least 99% identical to the nasal specimen. This represents the virus seed stocks.

### Virus Working Stocks

Viral working stocks were produced by infecting approximately 80% confluent T-150 flasks of VeroE6-TMPRSS2 cells at an MOI of 0.01 TCID50/cell using the seed stocks. Flasks were incubated for 1 hour at 33°C, replenished with fresh IM and incubated at 33°C until 75% CPE was observed. Supernatant was harvested, centrifuged at 400g for 10mins to remove cellular debris and aliquoted for storage at −65°C as working stocks.

### Amplicon-Based Whole Genome Sequencing and Lineage/Clade Designation

Specimen preparation, extractions, and whole genome sequencing were performed as described previously [40, 41]. NEBNext^®^ ARTIC SARS-CoV-2 Companion Kit (VarSkip Short SARS-CoV-2 # E7660-L) was used for library preparation and sequencing using the Nanopore GridION. Base-calling of reads was conducted using the MinKNOW, followed by demultiplexing with guppybarcoder requiring barcodes at both ends. Artic-ncov2019 medaka protocol was used for alignment and variant calling. Clades were determined using Nextclade beta v 1.13.2 (clades.nextstrain.org), and lineages were determined with Pangolin COVID-19 Lineage Assigner [42]. Sequences with coverage >90% and mean depth >100 were submitted to GISAID database. All clinical isolates contain the same mutations as were identified in the nasal specimen.

### Tissue Culture Infectious Dose 50 (TCID_50_)

Throughout this study, SARS-CoV-2 virus titers were determined using TCID_50_ described similarly in previous studies [43, 44]. 96-well plates were seeded with VeroE6-TMPRSS2 cells to achieve 80% confluency. Cells were washed with PBS+/+ (1X PBS supplemented with Ca^2+^/Mg^2+^) once and replenished with 180μL IM prior to infection. Viral samples were serially diluted in ten-folds, and 20μL of each serially diluted samples were incubated as sextuplicate on 96-well plates for five days. Cells were fixed with 4% formaldehyde overnight and stained with Napthol Blue Black to visually score CPE and calculate TCID_50_ using the Reed-Muench method [45].

### Methylcellulose Plaque Assay

VeroE6 or VeroE6-TMPRSS2 cells were seeded in 6-well plates to achieve 100% confluency. Virus stocks were serially diluted in ten-folds and 250μL of each dilutions were transferred into each well. Virus dilutions were removed after 1 hour incubation at either 33°C or 37°C, and 2ml of 1% methylcellulose overlay media were added to each well. To achieve distinguishable and visible plaques, cells were incubated for either 96 hours at 37°C or for 120 hours at 33°C when grown on VeroE6 cells and either for 48 hours at 37°C or for 72 hours at 33°C when grown on VeroE6-TMPRSS2 cells. Wells were fixed with 4% formaldehyde overnight and stained with Napthol Blue Black to count plaques. Plaque areas were determined by imaging wells and a standard ruler using a Nikon Fluorescence Dissection Microscope with an Olympus DP-70 camera. Images of wells were transferred to ImageJ software and a pixel-to-mm scale was established using the ruler. Plaques were traced and area (in mm^2^) was determined based on selected pixels. Plaque sizes were graphed using GraphPad Prism 9 software and ordinary one-way ANOVA with Tukey’s multiple comparison test was used for statistical analyses.

### Plasmas and Plaque Reduction Neutralization Test (PRNT)

CCP samples were collected between May 2020 and Mar 2021 [46]. As described in a previous study [47], VeroE6-TMPRSS2 cells were seeded and grown to 100% confluency in 6-well plates. Heat-inactivated CCP plasma samples were primarily diluted to 1:20 and serially diluted in two-folds until 1:2560. Each dilution and a non-plasma control were incubated with a SARS-CoV-2 isolate containing 100 PFU per experiment. Virus-plasma mixtures were incubated for 1 hour at room temperature and 250μL of the mixture was each overlaid on cell monolayers as duplicates for 1 hour at 37°C. Virus-plasma mixture were aspirated and 2ml of 1% methylcellulose (combination of 2% methylcellulose (Sigma, Cat#435244-250G) and equal parts of 2X MEM (Gibco, Cat#11935046) supplemented with 1% Penicillin and Streptomycin, 1% GlutaMAX and 10% FBS) were overlaid into each well for 48 hours at 37°C. Cells were fixed with 4% formaldehyde overnight and stained with Napthol Blue Black. IC_50_ was calculated based on PFU counts using the inhibition dose-response, non-linear regression model on GraphPad Prism 9 software.

### Low MOI Growth Curve Infections

For VeroE6-TMPRSS2 cell infections, VeroE6-TMPRSS2 cells were seeded onto 24-well plates to reach 100% confluency. Cells were washed with IM and infected with 100μL of diluted virus stocks at 0.01 MOI (TCID_50_/cells). Infections were done in four replicate wells per virus and incubated at either 33°C or 37°C for 1 hour. Virus inoculums were removed, and cells were washed and replenished with IM. At every 2-, 12-, 24-, 36-, 48-, 72-, 96- and 120-hours post infection, the supernatants were collected for viral titration. Microscope images were also taken off the first replicate well of each virus at all timepoints before collecting the supernatant using a EVOS AMEX1000 brightfield microscope using a EVOS AMEP 4632 4X/0.13 objective or EVOS AMEP 4634 20X/0.4 objective.

For hNEC infections, fully differentiated hNECs were acclimated at either 33°C or 37°C for at least 4 hours as similarly described in a different study [48]. hNECs were washed on the apical side with IM and the basolateral side was replenished with previously described ALI basal media and supplements. 100μL of virus dilutions to achieve a 0.1 MOI (TCID_50_/cells) infection was transferred to the apical side of each well. Infections were also done in quadruplicates per virus for each temperature. After 2-hour incubations at either 33°C or 37°C, cells were washed three times with PBS-/-(1X PBS without Ca^2+^/Mg^2+^) and left in ALI with basolateral media. At 2-, 12-, 24-, 36-, 48-, 72-, 96-, 120-, 144-, 168-hours post infection, IM was added for 10 minutes and collected for the apical supernatant while the basolateral media was replaced every 48 hours.

Infectious virus in all supernatants were quantified using TCID_50_ and graphed on GraphPad Prism 9 software. Two-way ANOVA with Tukey’s multiple comparison test was used to conduct statistical analyses.

### Immunofluorescence Microscopy

VeroE6-TMPRSS2 cells were grown on 8-chamber culture slides and infected with the SARS-CoV-2 isolates at an MOI of 1 at 37°C for 24 hours. The cells were washed with PBS, fixed using 4% paraformaldehyde for 10 minutes at room temperature, and permeabilized and blocked with PBS containing 0.5% Triton X-100 and 5% BSA. The cells were incubated with primary antibody diluted in PBS with 5% BSA that were specific to SARS-CoV-2 Spike protein (1:100; SinoBiological, Cat# 40590-D001). Fluorescently labeled secondary antibody AF488 (1:500 with final concentration of 4μg/ml; Thermofisher, Cat#A11013) was also diluted in PBS with 5% BSA. The slides were mounted on microscope slides using Prolong glass antifade mounting media containing Hoechst dye. Images were acquired using a brightfield Zeiss Axio Imager microscope with Zeiss-Plan Neofluar 40x/0.75 air objective and processed in ImageJ.

### Western Blotting for SARS-CoV-2 Spike Expression

VeroE6-TMPRSS2 cells seeded on 24-well plates were also infected with each isolates at an MOI of 1. Cell lysates were collected after 24 hours post-infection using RIPA lysis buffer (Sigma, Cat#20-188) supplemented with protease inhibitor (ThermoFisher, Cat#78430). Cells were sonicated in ice for 30 minutes and then pelleted by centrifuging at 12,000g for 12 minutes at 4°C. BCA protein assay kit (Pierce, Cat#23227) was used to determine the lysates’ protein concentration and 15ug of samples diluted to 37.5uL with PBS were then mixed with 12.5ul 4X Laemmli sample buffer (BioRad, Cat# 161-0747) and DTT (Pierce, Cat#20291) with a final concentration of 50mM, and boiled for 5 minutes and 30 seconds. Samples were loaded onto a 4-15% SDS-PAGE gel for separation and transferred to a PVDF membrane. The membranes were blocked with 5% milk-TBS (Tris-Buffered Saline) solution and immunoblotted using primary anti-S2 human/mouse chimera antibody (1:1000; SinoBiological, Cat#40590-D001) and anti-human AF488 (1:1000 with final concentration of 2μg/ml; ThermoFisher Cat#A-11013) for secondary conjugation. The membranes were also immunoblotted for house-keeping gene β-tubulin using primary rabbit anti-β-tubulin (1:2500 with final concentration 0.036 μg/ml; ThermoFisher, Cat#PA5-16863) and anti-rabbit AF647 (1:1000 with final concentration of 2μg/ml; ThermoFisher, Cat#A-31573) for secondary conjugation. Membrane images were taken using FluorChem Q and processed in ImageJ to quantify band intensity for protein expression levels. Cleavage efficiency was calculated by determining S2 subunit protein expression over total spike protein expression (full length spike and S2).

## Acknowledgments

We thank the members of the Pekosz laboratory and the laboratories of Sabra Klein and Kimberly Davis for helpful discussions and input on our experiments. This work was supported by the Johns Hopkins Center of Excellence in Influenza Research and Response (N2722014000007C and N7593021C00045) and the Richard Eliasberg Family Foundation.

